# Importins recognize the winged-helix fold of ETS transcription factors to mediate nuclear import

**DOI:** 10.64898/2026.01.06.697953

**Authors:** Michael McConville, Kaylee Lankford, Natalia Bernardes, Abby Walterscheid, Catherine Valadez, Ashley Brower, Yuh Min Chook, Glen Liszczak

## Abstract

Protein trafficking between the cytoplasm and the nucleus is a fundamental process in eukaryotic cell biology. While linear nuclear localization signals (NLSs) are well-characterized, many nuclear proteins lack a predictable NLS. Here, we identify the ETS domain, a DNA-binding winged-helix fold, from ETS family transcription factors as a structure-encoded NLS. We show that ETS domains mediate nuclear import through direct recognition by multiple nuclear transport receptors, including IPO9. Cryo-electron microscopy analysis of the EHF:IPO9 complex reveals that the IPO9 wraps around the ETS domain and engages structural features throughout the winged-helix fold. Biochemical studies demonstrate that the ETS domain DNA-binding helix is critical for importin recognition and for NLS activity in mammalian cells. Comparison of IPO9 bound to EHF and the histone H2A:H2B dimer reveals distinct interaction hotspots, illustrating how IPO9 employs unique combinatorial binding surfaces to accommodate structurally diverse cargos. These findings define a new class of globular NLSs and highlight the adaptability of importins in recognizing distinct protein folds.

**Significance Statement:** Nuclear import is essential for transcription factor function. However, many nuclear proteins lack recognizable nuclear localization signals (NLSs), leaving their trafficking mechanisms unresolved. Here, we identify the winged-helix DNA-binding domain of ETS transcription factors as a structure-encoded NLS shared across the ETS family of proteins. We show that multiple importins directly interact with this globular domain and define the molecular basis for cargo recognition by determining the cryo-EM structure of an importin bound to an ETS family protein. These studies establish a new class of globular NLSs and shed light on how individual importins can recognize diverse protein folds. We also provide mechanistic insight into nuclear trafficking defects that are caused by disease-linked ETS transcription factor mutations.

## Introduction

Mammalian cells employ diverse mechanisms to direct proteins to specific subcellular locations. These trafficking pathways regulate protein activity with high spatial and temporal precision. This is especially evident for nuclear proteins, which are translated in the cytoplasm and trafficked through the nuclear pore complex by members of the karyopherin protein family (1, 2). Karyopherin nuclear transport receptors include multiple importins and exportins with distinct cargo recognition properties. While many cargos use well-defined linear nuclear translocation signals to engage their cognate importin, there are also karyopherin-dependent trafficking mechanisms that do not operate through predictable signal sequences and remain poorly understood (1, 3–6). Notably, disease-associated mutations in nuclear proteins frequently coincide with subcellular mislocalization, although the mechanistic basis underlying such trafficking defects is often unclear (7–9). These observations highlight the need to elucidate karyopherin:cargo recognition mechanisms and to better understand this fundamental aspect of cellular signaling in both physiology and disease.

The known mechanisms by which importins recognize their cargo proteins are highly diverse (1). Among well-defined NLS motifs are: (i) the classical-NLS (cNLS) that is recognized by Importin subunit α (10), (ii) the proline-tyrosine (PY)-NLS that binds Transportin-1 (also known as Karyopherin-β2) (11), (iii) the isoleucine-lysine (IK)-NLS that is recognized by yeast Kap121 and its homolog Importin-5 (IPO5) (12), and (iv) the arginine-serine or arginine-serine-tyrosine (RS or RSY)-NLS that binds Transportin-3 (TPNO3) (13). Beyond these motifs, many proteins carry uncharacterized NLSs embedded within intrinsically disordered regions, while others rely on nuclear targeting elements located within structured domains.

All ten importins and three biportins have been implicated in recognizing stably folded protein domains in addition to or instead of linear NLS motifs (1, 14). However, relatively few structural studies have characterized importins bound to these ‘globular NLSs’. Notable examples of structure-encoded NLSs that bind importins include IMPβ with SREBP-2 (15) and SNAI1 (16), IPO4 with the H3-H4:ASF1 complex (17), IPO9 with the H2A:H2B dimer (18), TPNO3 with ASF/SF2 (19), and IPO13 with the Mago:Y14 dimer (20) and Ubc9 (21). These cargos are structurally diverse, and many can interact with multiple importin proteins (22). Moreover, individual importins can engage multiple distinct protein folds (14, 23). As a result, the molecular details that define nuclear import cargo recognition, especially globular NLS cargos, remain poorly understood and are not reliably predictable by computational approaches.

Previous studies have shown that importins can serve dual roles as nuclear transport receptors and chaperones for highly basic, aggregation-prone proteins, such as histones, ribosomal proteins, and various other DNA- and RNA-binding proteins (14, 17, 18, 23–27). Importantly, disease-associated mutations in transcription factors often lead to subcellular mislocalization (7, 8). For example, mutations throughout the ETS domain of the transcriptional repressor ETV6, which is a DNA-binding winged-helix fold that is common to all ETS family proteins, impair both DNA binding and nuclear localization (28–32). These mutations are strongly associated with familial thrombocytopenia and hematologic malignancies (32, 33). Notably, the ETS domain is also required for ETV6 nuclear localization, despite the absence of a predictable NLS motif (34). We therefore hypothesized that the ETV6 ETS domain may serve as a globular NLS. To better understand the cellular factors that govern ETV6 subcellular localization, we determined the nuclear import mechanism for this critical hematopoiesis regulator.

Here, we show that the ETS domain from ETS family transcription factors, including ETV6, functions as a globular NLS that is recognized by multiple importin proteins, including IMPβ1, IPO4, and IPO9. We determined the cryo-EM structure of IPO9 bound to the ETS protein EHF, revealing that IPO9 employs multiple HEAT repeats and intervening loops to envelop the winged-helix fold. Mutations within key IPO9 cargo-binding loops, as well as within the DNA-binding helix of EHF and ETV6 ETS domains, abolish the interaction. Comparisons of the IPO9:EHF structure with those of IPO9:RanGTP and IPO9:H2A:H2B define the molecular basis for RanGTP competition and reveal IPO9 cargo-specific binding mechanisms. These structural analyses underscore key features of IPO9 plasticity, including a multitude of cargo interfaces and conformational adaptability, that enable recognition of diverse folds.

## Results

### The ETS domain of ETS transcription factors functions as a nuclear localization signal

Consistent with previous studies (34), we found that deletion of the ETV6 C-terminus (residues 334-452) disrupted nuclear localization (Fig. S1). This region contains the winged-helix fold that mediates DNA binding (ETS domain; residues 334-430) and a short C-terminal inhibitory domain (CID; residues 421-452; Fig. 1a) (35). The CID is known to form an acidic α-helix that interacts with the ‘DNA-binding helix’ in the ETS domain to attenuate DNA binding activity (35–37). To further investigate the ETS-CID region as a potential atypical NLS, we employed a confocal microscopy-based cellular assay for NLS activity (Fig. 1b). In this assay, NLS activity is quantified by the nuclear accumulation of an otherwise cytosolic GFP-β-galactosidase (GFP-β-Gal) protein when fused to an NLS of interest.

**Fig. 1.**
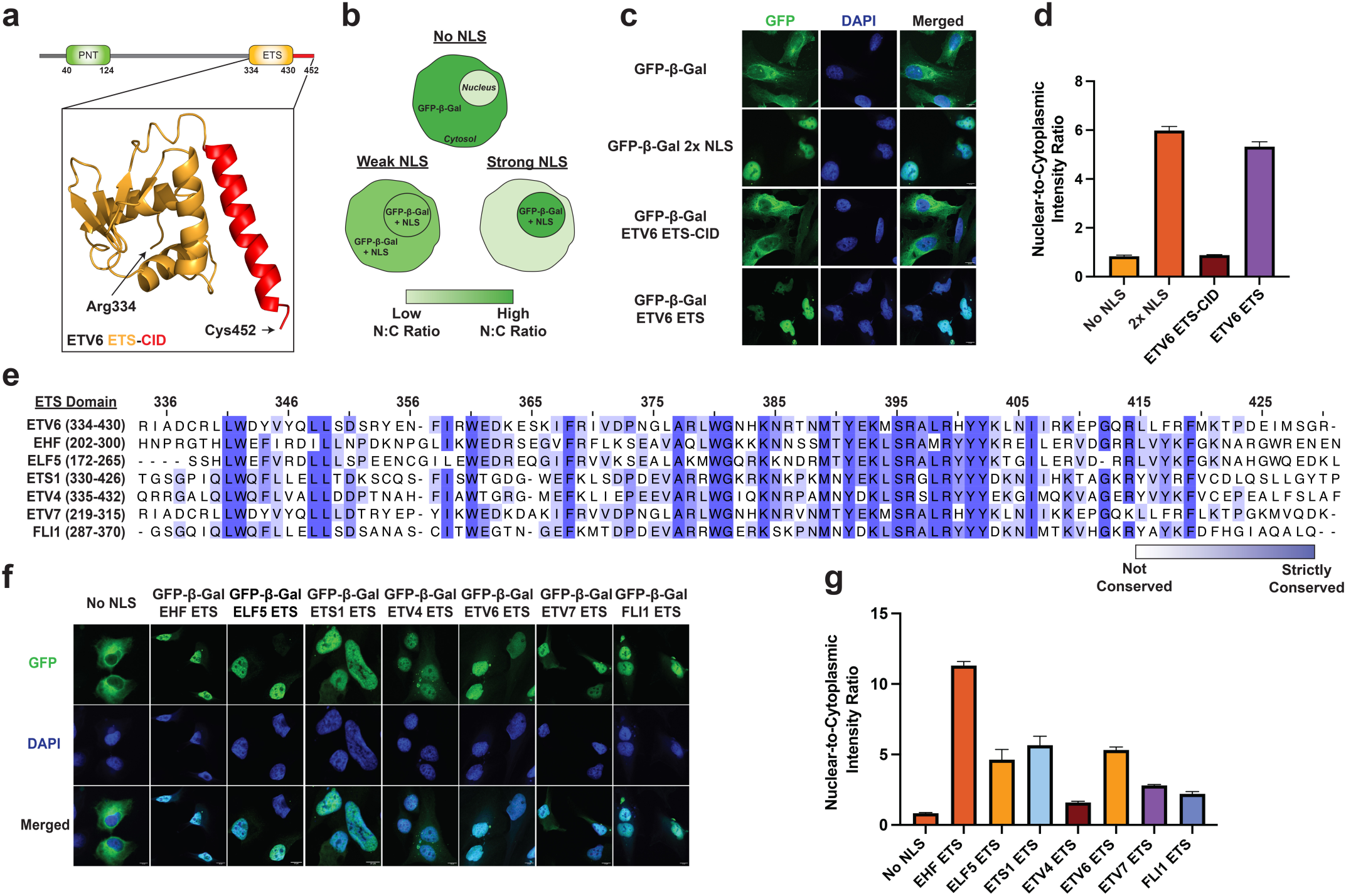
The globular ETS domain is a nuclear localization signal. a. Domain organization of ETV6 with a zoomed view of the ETS-CID fragment structure (orange and red, respectively). b. Schematic of the cellular NLS reporter assay. c. Fluorescence microscopy z-stack images of HeLa cells bearing the GFP-β-Gal NLS reporter fused to the SV40 large T antigen NLS or the indicated ETV6 fragment. Scale bar = 10 μm. d. Nuclear-to-cytoplasmic intensity ratio of the indicated NLS reporter protein shown in (c). Quantification was performed using fluorescence microscopy. Fluorescence intensity was determined for n = 3 viewing fields with greater than 50 cells per field. e. A multiple sequence alignment of the ETS domain from various ETS family transcription factors. f. Fluorescence microscopy z-stack images of HeLa cells bearing the GFP-β-Gal NLS reporter fused to the indicated ETS domain. Scale bar = 10 μm. g. Nuclear-to-cytoplasmic intensity ratio of the indicated reporter protein shown in (f). Quantification was performed using fluorescence microscopy. Fluorescence intensity was determined for n = 3 viewing fields with greater than 50 cells per field. ETV6 ETS domain N:C ratio calculation identical to panel (d).

Ectopic GFP-β-Gal-fusion constructs were stably integrated into HeLa cells. Cells expressing the exogenous fluorescent protein were enriched via FACS and analyzed by confocal microscopy (Fig. 1c). As expected, the unmodified GFP-β-Gal protein was predominantly cytoplasmic, with a nuclear-to-cytoplasmic fluorescence intensity ratio (N:C ratio) of 0.84 (Fig. 1d). Fusion of a known, potent NLS (2x-PKKKRKV from SV40 large T antigen (10)) to the C-terminus of GFP-β-Gal led to strong nuclear accumulation of the protein, yielding an N:C ratio of 6.0. Interestingly, fusion of the ETS-CID fragment had no detectable impact on protein localization, resulting in an N:C ratio of 0.89, comparable to that of the reporter protein lacking an NLS. Previous studies have shown that the ETS:CID interaction is considerably more stable in the isolated ETS-CID fragment than in full-length ETV6 (36). We therefore hypothesized that the CID may mask the NLS activity of the ETS domain in our assay. Remarkably, removal of the CID resulted in strong NLS activity, as the GFP-β-Gal-ETS domain fusion exhibited an N:C ratio of 5.3. We note that NLS activity has not been previously observed for a winged-helix or structurally similar fold, and the cognate nuclear import receptor is not known.

To further investigate the ETS domain as an atypical NLS, we generated stable cell lines expressing GFP-β-Gal fused to six distinct ETS family winged-helix DNA-binding domains (EHF, ELF5, ETS1, ETV4, ETV7, and FLI1). Domain boundaries were defined by alignment to the ETV6 ETS domain, with strict sequence conservation ranging from 39-80% (Fig. 1e). All six ETS domains exhibited an increased N:C ratio compared to the no NLS control, with EHF producing the strongest nuclear enrichment (N:C ratio = 11; Fig. 1f and g). Collectively, these results demonstrate that the winged-helix domain of multiple ETS family transcription factors can function as a globular NLS.

### Multiple importins directly engage ETS domains

Unlike cNLS motifs, globular domain NLSs and their cognate importins cannot be readily predicted by computational methods. To identify the nuclear import receptors that engage ETS domains, nine recombinant GST-tagged importins were purified to homogeneity and immobilized on glutathione agarose for immunoprecipitation analysis. Immobilized importins were incubated with the recombinant ETV6 ETS domain, washed thoroughly, and eluates were analyzed via SDS-PAGE to assess complex formation (Fig. 2a). We found that seven of the nine importins tested bound to the ETV6 ETS domain, with IMPβ, IPO4, and IPO9 exhibiting the most robust pull-down signals.

**Fig. 2.**
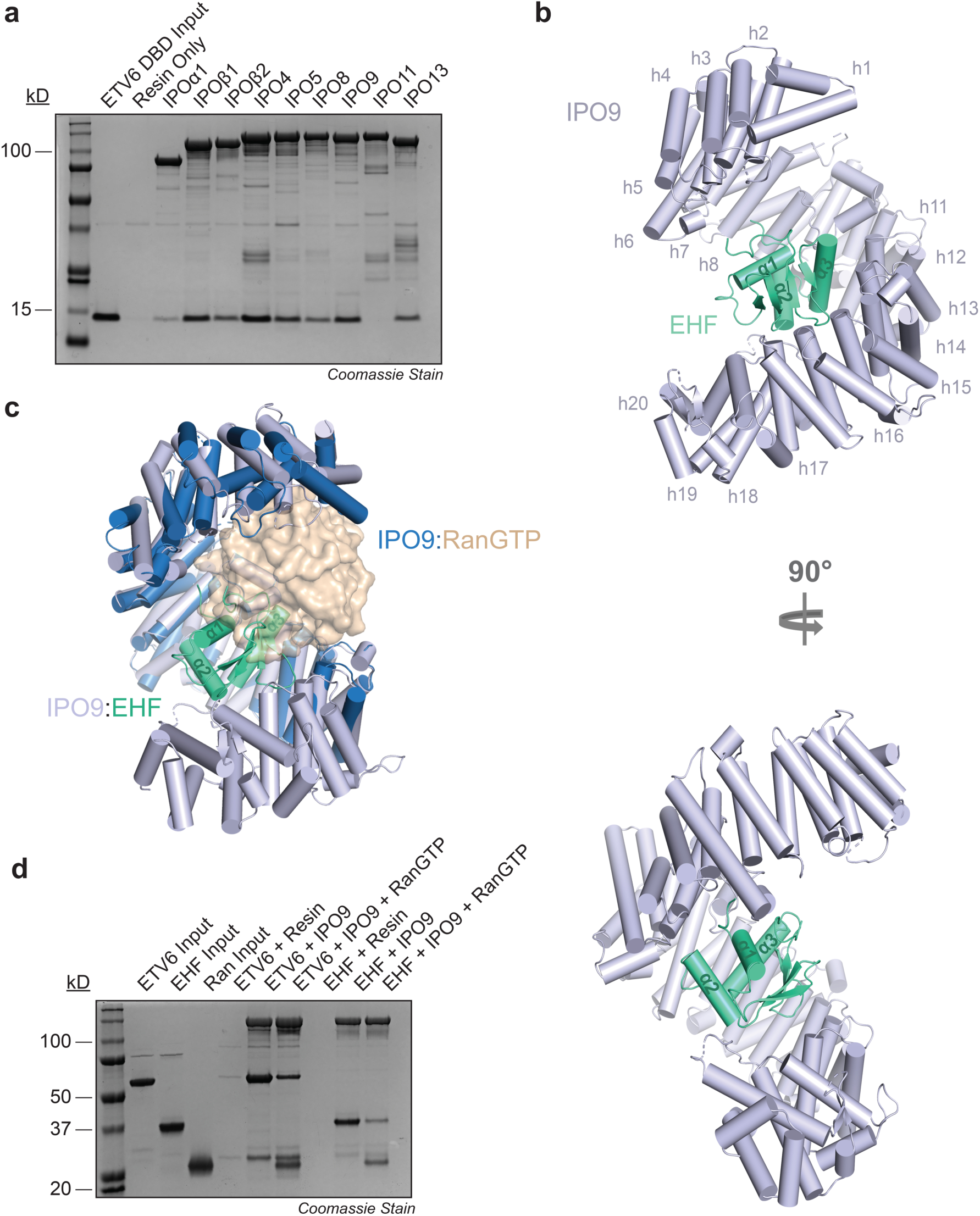
IPO9 engages the globular ETS domain. a. SDS-PAGE analysis of GST pull-down assays assessing the interaction between immobilized recombinant importin proteins (bait) and the recombinant ETV6 ETS domain (prey). b. Structural model of the IPO9:EHF complex as determined by cryo-EM. c. Structural overlay of the IPO9:RanGTP and IPO9:EHF complexes. d. SDS-PAGE analysis of GST pull-down assays assaying the effect of recombinant RanGTP on the interaction between immobilized GST-IPO9 (bait) and full-length ETV6 and EHF.

Comparable results were obtained using full-length recombinant ETV6 as prey, suggesting that importins can engage the ETS domain in the context of the intact protein (Fig. S2). Our findings align with multiple previous reports demonstrating that importins display overlapping cargo preferences, particularly for basic nucleic acid-binding domains (14, 17, 18, 22–27).

Having established that multiple importins bind the ETS domain, we next asked which importin:ETS domain complex would form the most robust complex for structural characterization. To determine this, we selected IMPβ, IPO4, and IPO9 for follow-up complex formation studies with the ETS domain from ETV6 and EHF, which exhibited the strongest NLS activity in our mammalian cell assay. A series of gel filtration chromatography experiments showed that while most ETS protein:importin complexes partially or completely dissociated during the run, the IPO9:EHF complex remained intact (Fig. S3). We therefore pursued cryo-EM structural analysis of IPO9 bound to full-length EHF.

### IPO9 wraps around the ETS winged-helix fold to orchestrate cargo recognition

We determined the cryo-EM structure of the IPO9:EHF complex at 3.48 Å resolution. Full-length IPO9 and the ETS domain of EHF (residues 204-298) were readily modeled into the electron density map (Fig. 2b and S4a), whereas no density was observed for the EHF N-terminal region (residues 1-203). Local resolution analysis indicated generally uniform map quality, with particularly well-resolved density at the IPO9:EHF interface (Fig. S4b and S4c). In contrast, lower local resolution was observed for the N-terminal HEAT repeats of IPO9, suggesting increased flexibility in this region when bound to EHF (Fig. S4b).

Structural analysis revealed that the twenty tandem HEAT repeats of IPO9 (h1-h20; each comprising α-helices A and B) form a superhelical architecture with an acidic concave surface, similar to that observed in the RanGTP- and H2A-H2B-bound IPO9 structures (Fig. 2b) (18, 38). The concave surface, including residues from multiple HEAT repeats and intervening loops, forms extensive contacts with the EHF ETS domain, burying 1,361.9 Å^2^ of the ETS domain surface area. Notably, structural alignment of IPO9:EHF and IPO9:RanGTP revealed substantial overlap between their binding sites (Fig. 2c). Furthermore, binding of both EHF and ETV6 to IPO9 was reduced in the presence of RanGTP in reconstituted protein pull-down assays (Fig. 2d). These results suggest that ETS domain cargo release in the nucleus follows the canonical RanGTP-dependent mechanism that has been established for most nuclear import cargos (3, 39).

Among the extensive interaction network, the α3 helix of EHF (EHF-α3) serves as a central recognition element and sits in a groove formed by three IPO9 helices, h11B, h12B, and h13B (Interface 1 in Fig. 3a). In ETS family winged-helices, the α3 helix is referred to as the ‘DNA-binding helix’ because it harbors multiple basic residues that intercalate DNA to mediate nucleic acid binding (Fig. 3b) (35). In ETV6, this same basic face of the α3 helix forms intramolecular contacts with the CID, explaining how DNA binding is autoinhibited in the intact protein (Fig. 3c) (37). Structural superposition of the ETV6 ETS-CID fragment with IPO9-bound EHF shows that the CID would sterically clash with IPO9 and clarifies why the ETV6 ETS-CID fragment fails to form a stable complex with importins (Fig. 3d and e). The basic face of the α3 helix, including K262, R265, R268, K272, and R273, forms electrostatic and hydrogen-bonding interactions with residues on IPO9 helices h11B, h13B, and a loop connecting h8A and h8B (h8^loop^; Fig. 3a). Additional EHF-α3-mediated contacts involve E261, Y269, Y270, which engage in a series of hydrophobic, electrostatic, and hydrogen-bonding interactions with residues in IPO9 helices h12B, h13B, and the h8^loop^.

**Fig. 3.**
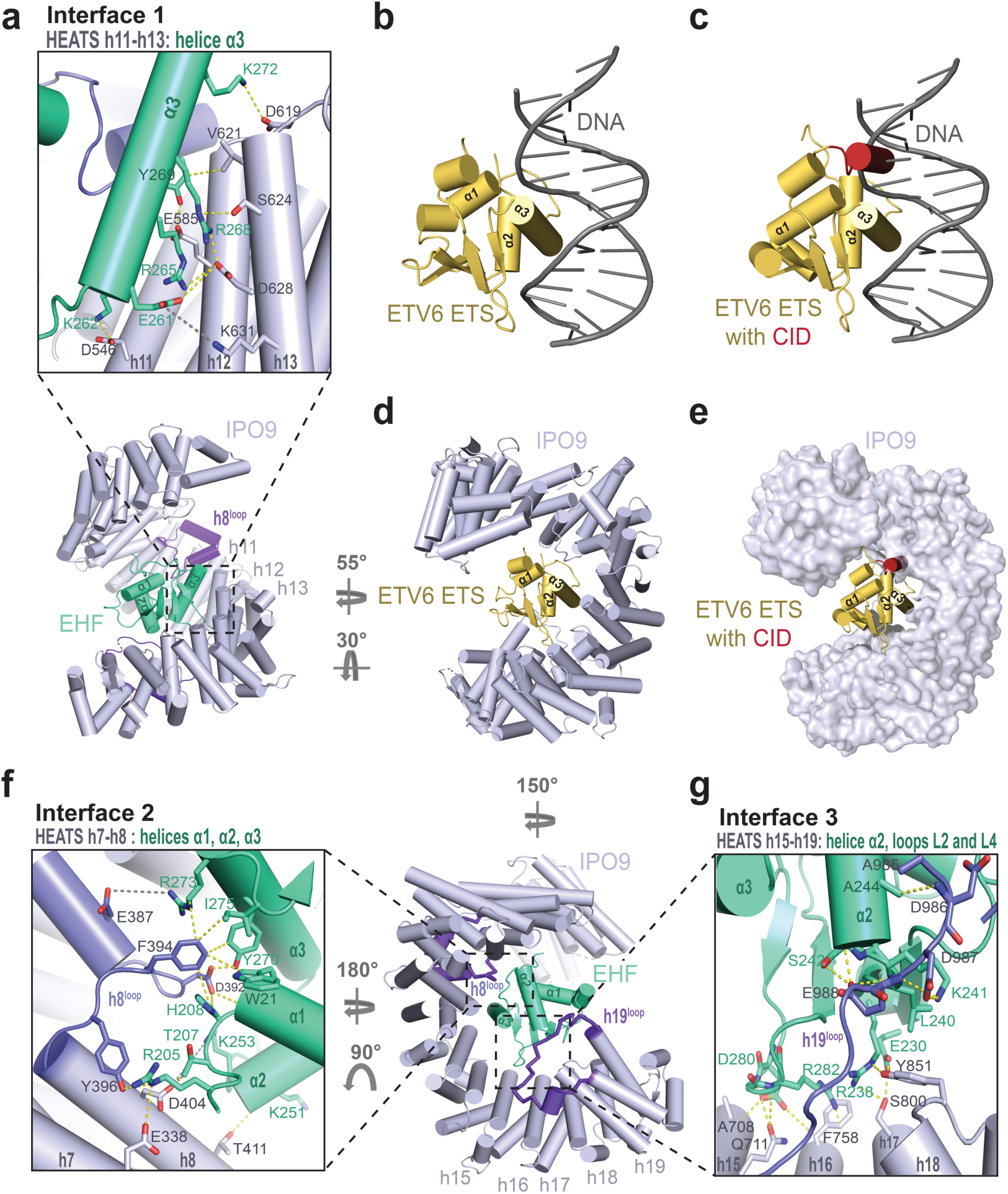
Multiple IPO9 interfaces contribute to ETS domain cargo recognition. a. Zoomed view of Interface 1. Residues involved in intermolecular interactions are indicated and shown in stick configuration. b. Structure of the ETV6 ETS domain bound to DNA from De, S., *et. al* (PBD ID: 4MHG) (35). c. Structural superposition of the ETV6 ETS-CID fragment from Coyne, H.J., *et. al* (PDB ID: 2LF7) (37) and the ETV6 ETS:DNA complex. d. Structural superposition of the ETV6 ETS domain with IPO9-bound EHF. EHF is omitted from the model for clarity. e. Structural superposition of the ETV6 ETS-CID fragment with IPO9-bound EHF. EHF is omitted from the model for clarity. f. Zoomed view of Interface 2. Residues involved in intermolecular interactions are indicated and shown in stick configuration. g. Zoomed view of Interface 3. Residues involved in intermolecular interactions are indicated and shown in stick configuration.

Beyond EHF-α3-dependent Interface 1, Interfaces 2, and 3 include amino acids spread out across various ETS domain secondary structure elements to interact with IPO9. Interface 2 employs side chains from EHF-α1 and the preceding loop to engage IPO9 residues in h7B, h8B, and the h8^loop^ (Fig. 3f). Interface 3 leverages residues in EHF-α2 and the central β-sheet to interact with IPO9 helices h15B, h16B, h17B, and the h19^loop^ (Fig. 3g). Both Interface 2 and Interface 3 include a distribution of hydrophobic, electrostatic, and hydrogen-bonding interactions.

### IPO9 exploits multiple substrate binding loops for ETS domain cargo recognition

To define the IPO9 surface elements that are required for ETS domain recognition, we purified a pilot series of IPO9 variants bearing single- or double-point mutations for immunoprecipitation analysis. However, none of these substitutions had a measurable effect on ETS domain binding in pull-down assays (Fig. S5). Given the extensive nature of the IPO9:ETS domain interface, we next generated four IPO9 variants containing clustered mutations targeted to Interface 1, 2, or 3. For Interface 1, we generated IPO9 variants in which either eight (IPO9-8x) or a more extensive set of twelve (IPO9-12x) residues were simultaneously mutated, all of which make contacts with EHF-α3. To probe Interfaces 2 and 3, the h8 loop (residues 371-396) or h19 loop (residue 941-995), respectively, was replaced with a glycine-serine-rich linker as previously described (Δh8^loop^ and Δh19^loop^) (18).

We found that Interface 1 required extensive mutagenesis to elicit a detectable decrease in ETS domain binding. The IPO9-8x variant showed no clear binding defect, whereas the more comprehensive IPO9-12x mutant displayed a modest reduction in binding to both EHF and ETV6 in recombinant protein immunoprecipitation assays (Fig. 4a and b). For Interfaces 2 and 3, the Δh8^loop^ and Δh19^loop^ variants each caused a modest decrease in ETS domain binding (Fig. 4a and b). However, combining these loop substitutions to create a Δh8^loop^/Δh19^loop^ variant resulted in an almost complete loss of binding with both EHF and ETV6. By comparison, this double loop mutant exhibited only a modest decrease in binding to RanGTP (Fig. S6), indicating that the ETS domain cargo is particularly dependent on these loops. Together, these results indicated that all three interface hotspots contribute to ETS domain cargo recognition.

**Fig. 4.**
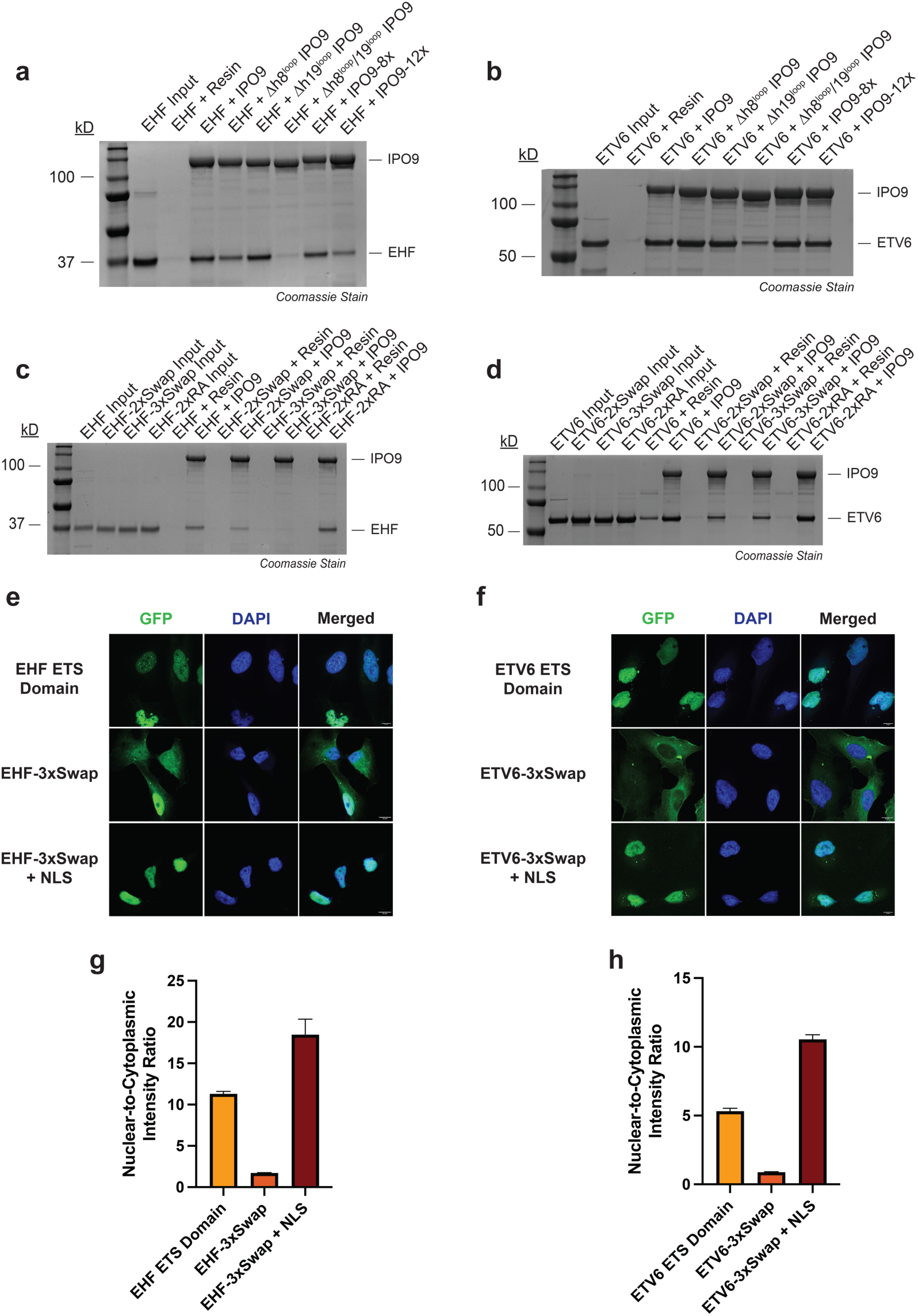
Distinct structural elements of IPO9 and the ETS domain cooperate to mediate structure-encoded NLS recognition and nuclear import activity. a. SDS-PAGE analysis of GST pull-down assays assessing the interaction between immobilized recombinant IPO9 protein variants (bait) and full-length, recombinant EHF (prey). ΔH8 = residues 371-396 replaced with glycine-serine linker; ΔH19 = residues 941-995 replaced with glycine-serine linker; IPO9-8x = E387K/D392K/F394A/D546K/E585K/D619K/D628K/K631E; IPO9-12x = IPO9-8x + F758A/Y851A/D987K/E988K. b. SDS-PAGE analysis of GST pull-down assays assessing the interaction between immobilized recombinant IPO9 protein variants (bait) and full-length, recombinant ETV6 (prey). IPO9 variants are defined in (a). c. SDS-PAGE analysis of GST pull-down assays assessing the interaction between immobilized recombinant IPO9 (bait) and the full-length, recombinant EHF protein variants (prey). EHF-2xSwap = R265E/R268E; EHF-3xSwap = K262E/R265E/R268E; EHF-2xRA = R265A/R268A. d. SDS-PAGE analysis of GST pull-down assays assessing the interaction between immobilized recombinant IPO9 (bait) and the full-length, recombinant ETV6 protein variants (prey). ETV6-2xSwap = R396E/R399E; ETV6-3xSwap = K393E/R396E/R399E; ETV6-2xRA = R396A/R399A. e. Fluorescence microscopy z-stack images of HeLa cells bearing the GFP-β-Gal NLS reporter fused to the indicated EHF fragment. Scale bar = 10 μm. f. Fluorescence microscopy z-stack images of HeLa cells bearing the GFP-β-Gal NLS reporter fused to the indicated ETV6 fragment. Scale bar = 10 μm. g. Nuclear-to-cytoplasmic intensity ratio of the indicated NLS reporter-EHF fusion protein shown in (e). Quantification was performed using fluorescence microscopy. Fluorescence intensity was determined for n = 3 viewing fields with greater than 50 cells per field. h. Nuclear-to-cytoplasmic intensity ratio of the indicated NLS reporter-ETV6 fusion protein shown in (f). Quantification was performed using fluorescence microscopy. Fluorescence intensity was determined for n = 3 viewing fields with greater than 50 cells per field. ETV6 ETS domain N:C ratio calculation identical to Fig. 1d.

### The ETS ‘DNA-binding helix’ is an essential feature for IPO9 recognition

As observed with IPO9, single ETS domain point mutations within Interface 1, 2, or 3 had no detectable effect on IPO9 binding in pull-down analyses (Fig. S7). Given that EHF-α3 (the DNA-binding helix) is intimately buried within the IPO9:EHF interface we focused on EHF-α3 for combinatorial mutation analysis. The K262, R265, and R268 residues of EHF (K393, R396, and R399 in ETV6, respectively) are key residues at the basic face of the DNA-binding helix that engage the IPO9 concave surface. We first generated an EHF R265A/R268A (EHF-2xRA) double mutant protein and a corresponding ETV6 R396A/R399A (EHF-2xRA) variant. Neither of these mutants appreciably altered IPO9 binding in immunoprecipitation assays (Fig. 4c and d). Given that these residues interact with complementary acidic side chains on IPO9, we next introduced charge-swap substitutions to create electrostatic repulsion at the interface. ETS domain charge-swap variants included EHF R265E/R268E (EHF-2xSwap), EHF K262E/R265E/R268E (EHF-3xSwap), and their synonymous ETV6 counterparts. In contrast to the 2xRA DNA-binding helix variants, all EHF- and ETV6-2xSwap and 3xSwap mutant proteins exhibited impaired binding to IPO9 in reconstituted protein immunoprecipitation analyses (Fig. 4c and d).

We next investigated the impact that DNA-binding helix charge swap mutations have on ETS domain nuclear import activity in a cellular context. We generated GFP-β-Gal constructs fused to the EHF-3xSwap and ETV6-3xSwap ETS domains for fluorescence microscopy-based subcellular localization analysis. Consistent with our findings in reconstituted protein pull-down assays, ETS domain charge-swap mutations severely impaired nuclear accumulation of the GFP-β-Gal reporter protein (Fig. 4e and f). The EHF-3xSwap and ETV6-3xSwap exhibited N:C ratios 6.6-fold and 6.1-fold less than their wild-type counterparts, respectively (Fig. 4g and h). In both cases, inclusion of the SV40-NLS at the C-terminus of the reporter construct rescued nuclear localization of the GFP-β-Gal-3xSwap proteins, strongly suggesting cytoplasmic enrichment of the 3xSwap-containing reporters is due to disruption of ETS domain:importin interactions in the cellular environment.

### IPO9 recognizes diverse cargo folds through unique combinations of interaction hotspots

Prior to this work, the histone H2A-H2B heterodimer was the only folded domain cargo for which an IPO9-bound structure had been determined (18). However, the H2A-H2B dimer does not assume a winged-helix-like fold and, unlike ETS domains, RanGTP is not sufficient to release the H2A-H2B complex from IPO9. A comparison of the EHF- and H2A-H2B-bound structures revealed that very few IPO9 interface residues are utilized for both cargos, and the overall cargo binding topologies are remarkably different (Fig. 5a). While the EHF ETS domain is buried relatively deep into the concave pocket of the IPO9 superhelical fold, the H2A-H2B heterodimer is clamped by the HEAT repeats at the two termini of IPO9 (Fig. 5b).

**Fig. 5.**
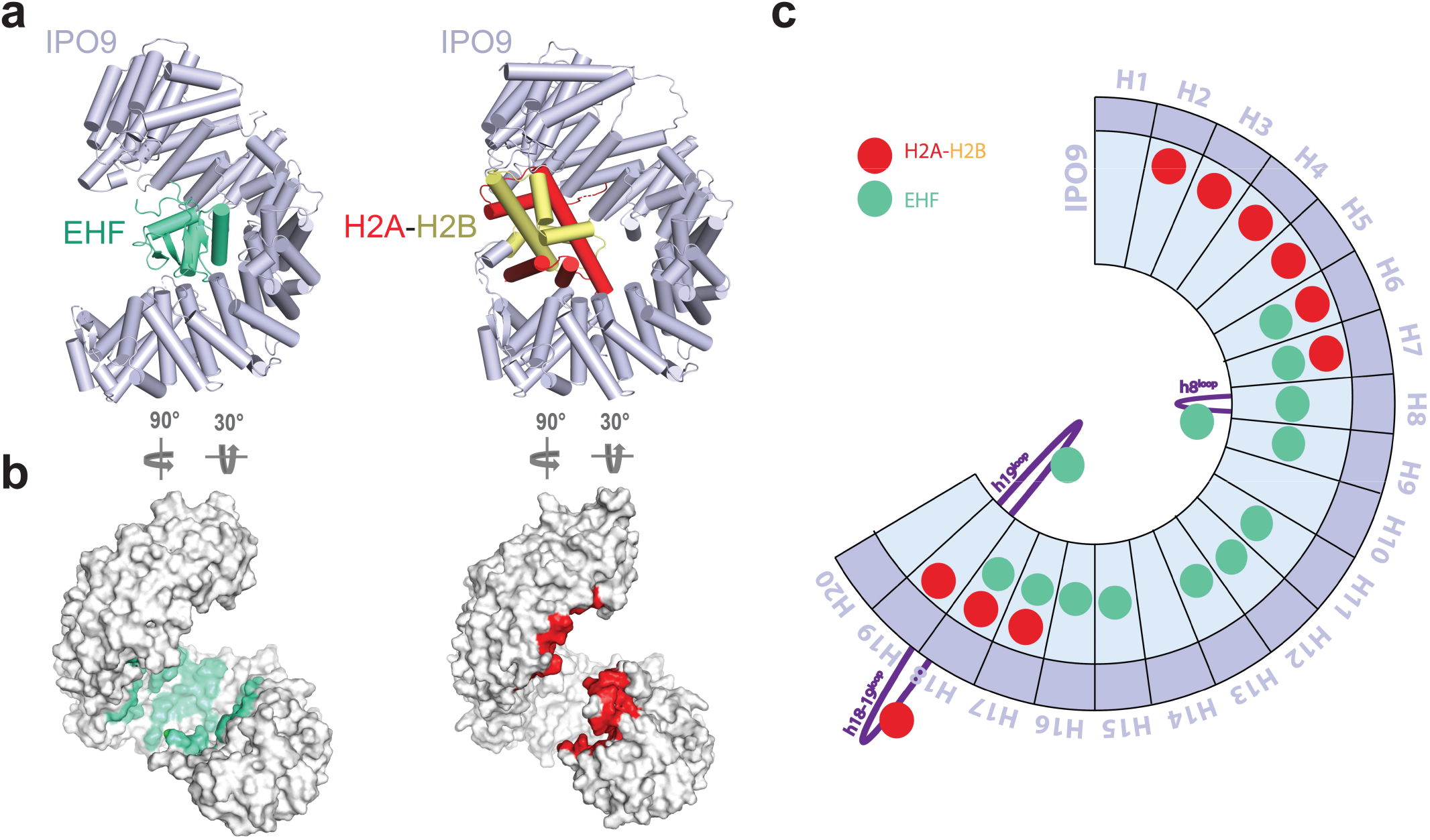
IPO9 employs unique cargo recognition mechanisms to accommodate the ETS domain and the H2A:H2B dimer. a. Side-by-side structural comparison of the IPO9:EHF and IPO9:H2A-H2B complexes. b. Surface representation of IPO9 with cargo contact interfaces highlighted (EHF in green and H2A-H2B in red). c. Schematic representation of HEAT repeats in IPO9 with contact sites for EHF and H2A-H2B shown as green and red circles, respectively. Note: H2A-H2B disordered tail contact sites do not contribute to IPO9 binding affinity and have been omitted here to emphasize critical interaction hotspots (18).

The ETS domain cargo exploits two IPO9 features that were not observed in the H2A-H2B study (Fig. 5c and d). First, the DNA-binding helix that we have identified as critical for importin recognition sits in a groove formed by IPO9 HEAT repeats, none of which make direct contacts with H2A-H2B. Second, the IPO9 h19 loop, which is critical for ETS domain recognition according to our biochemical analyses, could not be modeled in the H2A-H2B:IPO9 structure due to missing density. Inversely, IPO9 HEAT repeats 2-5 all form contacts with the H2A-H2B cargo while making no contacts with the ETS domain. The interfaces overlap at IPO9 HEAT repeats 7 and 8, as well as the h8^loop^, which contact a short segment of the H2B N-terminal tail. However, this interface contributes little to histone binding, as removal of either the H2B tail or the IPO9 h8^loop^ does not affect binding affinity of H2A-H2B for IPO9 (18).

## Discussion

Here, we found that multiple importin proteins can interact with the winged-helix fold from ETS family transcription factors to mediate nuclear import. All seven ETS domains tested in our study exhibited NLS activity in mammalian cells, albeit with varying strengths. These differences may reflect variations in the intrinsic stability of each isolated domain or the evolution of distinct importin-binding affinities that allow for regulated nuclear import in different biological contexts. Of these seven ETS proteins analyzed, six (EHF, ELF5, ETV4, ETV6, ETV7, and FLI1) lack a predictable cNLS based on the commonly used NLStradamus server (40). We therefore propose that the winged-helix domain functions as a structure-encoded NLS in many ETS family proteins. It is not yet clear if this property will extend to other nuclear proteins containing a winged-helix domain. This will likely be strongly influenced by electrostatic surface features of each fold and by the influence of neighboring domains or binding partners.

Previous work from our lab and others has shown that point mutations within the ETV6 ETS domain, including L349P, R369Q, R399C, and R418G, cause ETV6 mislocalization in cells (28–32). Among these, R369Q and R399C are known to disrupt DNA binding in reconstituted protein assays (30). In this study, we found that the ETV6 L349P and R418G protein variants fail to yield soluble protein when expressed in *E. coli*. This observation suggests that the L349P and R418G substitutions lead to misfolding of the winged-helix domain, thereby destroying the NLS and leading to cytoplasmic accumulation. In contrast, mutations at the R369 and R399 positions did not impair production of soluble recombinant protein or affect the ETV6:IPO9 interaction. Notably, soluble expression of full-length ETV6 in *E. coli* requires a point mutation in the N-terminal PNT oligomerization domain to block homo-oligomerization (41). It is therefore possible that the R369Q and R399C mutations may destabilize the protein only in the context of a construct bearing an oligomerization-competent PNT domain. It is also possible that these mutant proteins are less stable only in the mammalian cell environment, or that DNA-binding-deficient ETV6 is more readily exported out of the nucleus.

Comparisons between our experimentally determined EHF:IPO9 structure and the AlphaFold-multimer-predicted model reveal that, in its current state, AlphaFold Multimer V3 is unable to correctly predict key molecular details of the interaction interface (Fig. S8) (42). Thus, experimental approaches will be required to elucidate how other importins, such as IMPβ, recognize the winged-helix fold of ETS family proteins. Given that the 3xSwap mutations disrupt ETS domain NLS activity in cellular assays where multiple importin proteins exist, we expect that the DNA-binding helix forms critical interface contacts with other ETS domain binding-competent importins. Furthermore, our findings show that IPO9 exploits unique combinations of cargo binding sites to accommodate structurally diverse globular NLSs. This is further supported by structural work with the yeast homolog of IPO9, Kap114p, bound to the structured core domain of the yeast TATA-box binding protein (yTBP) (43). Indeed, the Kap114p:yTBP exhibits a cargo recognition mechanism that is unique from that of both EHF and H2A-H2B (Fig. S9a-c). Future studies are needed to further explore this principle in greater depth, particularly with IPO9 cargos that are structurally distinct from the winged-helix and H2A-H2B dimer folds.

## Materials and Methods

Recombinant ETS family transcription factors, importin proteins, and all variants thereof were cloned and expressed in *E. coli* and purified to homogeneity via optimized chromatographic workflows. Protein-protein interactions were assessed using GST pull-down assays with purified components. Structural studies were performed by assembling and purifying the IPO9:EHF complex, which was then applied to holey carbon grids and vitrified for cryo-EM analysis (Titan Krios). Cell-based localization experiments were conducted using lentiviral expression of ETS constructs in HeLa cells, followed by immunofluorescence staining, confocal microscopy analysis, and quantitative nuclear-to-cytoplasmic ratio analysis. For detailed Materials and Methods, see Supporting Information.

## Acknowledgements

We thank the UT Southwestern Structural Biology Laboratory for assistance with Cryo-EM data collection and processing and members of the Liszczak and Chook laboratories for productive discussions. This work was supported by grants from the Cancer Prevention and Research Institute of Texas (RR180051 to G.L.), the Welch Foundation (I-2039-20230405 to G.L., I-1532 to Y.M.C.), and the NIH (5F30HL167629 to M.M., 1R35GM147140 to G.L., R35GM144137 to Y.M.C.). G.L. is a Virginia Murchison Linthicum Scholar in Medical Research. Y.M.C. is the Alfred and Mabel Gilman Chair in Molecular Pharmacology and the Eugene McDermott Scholar in Biomedical Research.

