## Supporting Information for "Importins recognize the winged-helix fold of ETS transcription factors to mediate nuclear import"

**This PDF file includes**:

Supporting Information Materials and Methods

Figures S1-S9

Table S1

Supporting Information References

### Supporting Information Materials and Methods

#### Molecular cloning

All PCR amplification steps utilized the KAPA HiFi DNA Polymerase according to manufacturer's protocols. Inverse PCR was used to install single point mutations and Gibson Assembly with gene block gene fragments was used to install batch mutations. All DNA oligonucleotides were synthesized by Integrated DNA Technologies (Coralville, IA). Viral plasmid constructs were generated by restriction enzyme cloning. All plasmids from this study were whole plasmid sequenced by GENEWIZ (South Plainfield, NJ), Eurofins Genomics (Louisville, KY), or Plasmidsaurus (Louisville, KY). Plasmids that were used for recombinant protein production were transformed into Mach1 *E. coli* cells and plasmids that were used for Lentivirus production were transformed into Stbl3 *E. coli* cells.

#### Recombinant protein production

pGEX plasmids encoding GST-TEV-ETV6 protein (including wild-type, variants, full-length, and ETS domain fragment) were transformed into Rosetta2 (DE3) pLysS *E. coli* cells and inoculated in 6 L of Luria Broth (Miller). Note: to obtain soluble ETV6 protein, all full-length ETV6 *E. coli* expression constructs carry a V112E mutation that disrupts PNT domain oligomerization (1). Cells were grown in a shaker at 37°C to an $OD_{600}$ of 0.6 and protein expression was induced with 0.5 mM IPTG at 16°C overnight. Cells were sonicated in a lysis buffer that contained 50 mM HEPES (pH 7.5), 500 mM NaCl, 5 mM β-ME, and 1 mM PMSF. The soluble lysate was isolated by centrifugation at 40,000 RCF for 30 min at 4°C. The soluble fraction was incubated with GSH resin for 1 h at 4°C and the resin washed with 50 CV of lysis buffer without PMSF. The protein was eluted from resin in 30 mL of elution buffer containing 50 mM HEPES (pH 8.0), 100 mM NaCl, 2 mM DTT, and 20 mM glutathione. The GST tag was cleaved via incubation with TEV protease overnight at 4°C. The cleaved protein was diluted two-fold in 50 mM HEPES and the pH was adjusted to 6.5. Diluted protein was immobilized on a cation exchange column (HiTrap SP; Cytiva) that had been pre-equilibrated with a low salt buffer containing 50 mM HEPES (pH 6.5), 100 mM NaCl, and 2 mM DTT. The column was eluted with a linear gradient ranging from 100 mM NaCl to 1 M NaCl over 15 CV. Pure fractions as judged by SDS-PAGE analysis were pooled, NaCl concentration was adjusted to ~500 mM, and pH was adjusted to 7.5. Samples were concentrated using an Amicon Ultra Centrifugal filter (Millipore; 50 kDa MWCO) and glycerol was added to ~20%. Protein was aliquoted into single-use fractions, flash frozen, and stored at –80°C.

pET30 plasmids encoding 6xHis-SUMO-EHF (including wild-type, variants, full-length, and ETS domain fragment) were transformed into Rosetta2 (DE3) pLysS *E. coli* cells and inoculated in 6 L of Luria Broth (Miller). Cells were grown in a shaker at 37°C to an $OD_{600}$ of 0.6 and protein expression was induced with 0.5 mM IPTG at 16°C overnight. Cells were sonicated in a lysis buffer containing 50 mM HEPES (pH 7.5), 500 mM NaCl, 5 mM β-ME, and 1 mM PMSF. The soluble lysate was isolated by centrifugation at 40,000 RCF for 30 min at 4°C. The soluble fraction was incubated with Ni-NTA resin for 1 h at 4°C and the resin washed with 50 CV of lysis buffer supplemented with 25 mM imidazole. The protein was eluted from resin in 30 mL buffer containing 500 mM NaCl, 50 mM HEPES, 2 mM DTT, and 250 mM imidazole. The eluant was incubated with Ulp1 protease overnight at 4°C to remove the SUMO tag. Cleaved protein was diluted five-fold in 50 mM HEPES and pH was adjusted to 6.5. The diluted protein was applied to a cation exchange column (HiTrap SP; Cytiva) that had been pre-equilibrated with a low salt buffer containing 50 mM HEPES (pH 6.5), 100 mM NaCl, and 2 mM DTT. The column was eluted with a linear gradient from 100 mM NaCl to 1 M NaCl over 15 CV. Pure fractions as judged by SDS-PAGE analysis were pooled, NaCl concentration was adjusted to ~500 mM, and pH was adjusted to 7.5. Samples were concentrated using an Amicon Ultra Centrifugal filter (Millipore; 30 kDa MWCO) and glycerol was added to ~20%. Protein was aliquoted into single-use fractions, flash frozen, and stored at –80°C.

GST-tagged importin proteins (IPO4, IPO9, and KPNB1) were purified as previously described with several alterations (2, 3). Briefly, importin expression plasmids were transformed into Rosetta2 (DE3) pLysS *E. coli* cells and inoculated into 6 L of Terrific Broth (Fisher). Cells were grown in a shaker at 37°C to an $OD_{600}$ of 0.6 and protein expression was induced with 0.5 mM IPTG at 25°C for 10 h. Cells were sonicated in buffer containing 50 mM Tris-HCl (pH 7.5), 500 mM NaCl, 20 % glycerol, 2 mM DTT, and protease inhibitors (1 mM PMSF + Roche cOmplete protease inhibitor cocktail tablet). Soluble lysate was isolated by centrifugation at 40,000 RCF for 30 min at 4°C. The soluble fraction was incubated with GSH resin for 1 h at 4°C. The resin was first washed with 10 CV of lysis buffer supplemented with 2 mM EDTA. The second wash consisted of 10 CV with a lower salt buffer (Wash Buffer 2; 50 mM Tris-HCl (pH 7.5), 150 mM NaCl, 20% glycerol, and 2 mM $MgCl_2$). The third wash consisted of 10 CV of Wash Buffer 2 supplemented with 5 mM ATP. The fourth wash consisted of 10 CV of Wash Buffer 2. Lastly, protein was eluted in 30 mL of a buffer containing 50 mM Tris-HCl (pH 8.0), 150 mM NaCl, 20% glycerol, 2 mM $MgCl_2$, and 20 mM glutathione. The protein was immobilized on an anion exchange column (HiTrap Q; Cytiva) that had been pre-equilibrated with a low salt buffer containing 50 mM Tris-HCl (pH 7.5), 150 mM NaCl, 20% glycerol, and 2 mM DTT. The column was eluted with a linear gradient ranging from 150 mM NaCl to 1 M NaCl over 15 CV. Pure fractions as judged by SDS-PAGE analysis were pooled and samples were concentrated using an Amicon Ultra Centrifugal filter (Millipore; 50 kDa MWCO). Protein was aliquoted into single-use fractions, flash frozen, and stored at –80°C.

All other GST-tagged importin proteins in the recombinant importin protein panel and RanGTP were prepared exactly as previously described (4, 5).

**Multiple Sequence Alignment of ETS Family ETS Domains**

Protein sequence homology analysis for the ETS domain of ETS family proteins was analyzed by the online multiple sequence alignment tool (https://www.genome.jp/tools-bin/clustalw) (6). The final alignment image was generated and formatted using Jalview (7) and sequences were colored by percentage identity to show conservation.

**GST-tagged, recombinant importin:ETS protein pull-down assays**

Glutathione resin was pre-equilibrated in Immunoprecipitation (IP) Buffer (100 mM or 150 mM NaCl, 2 mM $Mg(OAc)_2$, 20 mM HEPES (pH 7.5), 1 mM EGTA, 15% glycerol, and 2 mM TCEP). Recombinant GST-importin proteins (60 µg) were incubated with 25 µl of glutathione resin (50 µl of slurry) at 4°C for 1 h. Following incubation, resin was washed three times with 500 µl of IP Buffer each. After the final wash, 60 µg of untagged recombinant ETS protein (ETV6 or EHF) was added to resin and the solution was brought to a final volume of 100 µl with IP Buffer. Proteins were incubated on resin at 4°C for 20 min.

Following incubation with ETS protein, the resin was washed three times with 500 µl of IP Buffer. After the final wash, resin was resuspended in 40 µl IP Buffer supplemented with 20 mM glutathione and incubated at 4°C for 20 min. Solutions were briefly spun to pellet resin and 30 µl of supernatant was collected. Supernatant was mixed with SDS loading dye, boiled, and analyzed by SDS-PAGE analysis. Coomassie staining was performed and gels were imaged on a Bio-Rad ChemiDoc.

*For RanGTP competition*: Following incubation with ETS protein, 60 µg of recombinant RanGTP was added directly to the resin and the volume was brought to 150 µl with IP Buffer. In control samples, the volume was brought to 150 µl with IP Buffer without addition of RanGTP. Samples were then incubated at 4°C for 20 min. Wash and elution steps were then performed as described above.

**IPO9:EHF complex assembly for structural analysis**

Full-length, untagged IPO9 and full-length, untagged EHF were incubated in a 1:5 molar ratio for 30 min on ice. The complex was then injected onto a gel filtration column (Superdex 200 Increase 10/300 GL; Cytiva) that had been pre-equilibrated with a buffer containing 20 mM HEPES (pH 7.4), 150 mM NaCl, 10% glycerol, 2 mM TCEP. Fractions containing the IPO9:EHF complex were identified via SDS-PAGE analysis, pooled, and concentrated to ~4 mg/mL total protein before preparing grids for data collection.

**Cryo-EM grid preparation and data collection**

Samples of the IPO9:EHF complex were applied to holey carbon grids (Quantifoil R1.2/1.3, 300-mesh copper) and vitrified using a Vitrobot Mark IV (Thermo Fisher Scientific). Prior to sample application, grids were glow-discharged in a PELCO easiGLOW unit (Ted Pella) for 60 s at 30 mA. One grid displaying optimal ice thickness and particle distribution was selected for data collection.

A 24 h data acquisition session was carried out at the Cryo-Electron Microscopy Facility at the University of Texas Southwestern (UTSW CEMF) using a Titan Krios microscope (Thermo Fisher Scientific) operated at 300 kV and equipped with a Gatan K3 direct electron detector. Data were collected in correlated double-sampling super-resolution mode at a nominal magnification of 105,000×, corresponding to a calibrated pixel size of 0.415 Å. A total of 5,338 movies was recorded using SerialEM (8), with a defocus range of –0.9 to –2.2 µm. Each movie consisted of 60 frames acquired over 5.4 s at an exposure rate of approximately 8 $e^{-}$/pixel/s.

**Cryo-EM data processing**

All image processing was performed in cryoSPARC (9). Movies were subjected to Patch Motion Correction and Patch CTF Estimation. An initial set of particles was extracted using Blob Picker and refined through three rounds of 2D classification. These particles served as templates for Template Picker, yielding a total of ~260,000 particles after three rounds of 2D classification. The selected particles were used to generate five *ab initio* models, followed by heterogeneous refinement. The best-resolved class, containing ~130,000 particles, was subjected to non-uniform refinement, resulting in a final reconstruction at 3.5 Å resolution.

**Cryo-EM model building, refinement, and analysis**

Atomic coordinates for IPO9 were taken from the crystal structure of the IPO9:H2A–H2B complex (PDB 6N1Z) (3). Coordinates for the EHF DNA-binding domain were obtained from the AlphaFold prediction of full-length EHF (AF-Q9NZC4-F1-v6) (10). These initial models were docked into the cryo-EM density maps using UCSF Chimera (11), followed by iterative model building in Coot (12) and refinement in Phenix (13). Final refinement steps included ISOLDE within UCSF ChimeraX (14). The refined structure was analyzed using CONTACT/ACT (15) and PDBe PISA (16) to identify intermolecular interactions and calculate solvent-accessible surface areas, respectively. PyMOL (17) was used for structural visualization and figure preparation. For data and model statistics, see Table S1.

**Cell culture**

HeLa cells and HEK293T cells were cultured in DMEM supplemented with 10% heat-inactivated FBS and 1% penicillin/streptomycin at 37°C and 5% $CO_2$.

**Lentivirus generation**

All lentivirus production was done using the HEK293T Lenti-X cell line, the pMD2.g lentivirus packaging plasmid, and the psPAX2 lentivirus packaging plasmid. To stably express constructs for localization studies, all gene inserts were PCR amplified and cloned into either the pHAGE-IRES-zsGreen vector, the pLVX-IRES-zsGreen vector using the BsmBI and BamHI restriction sites, or the pHAGE-GFP-β-Gal vector using a custom C-terminal MCS with BamHI and XhoI restriction sites.

On day 1, Lenti-X cells were seeded into 10-well plates. On day 2, each well was transfected with 2.5 µg pLVX or pHAGE expression plasmid, 1.5 µg psPAX2 and 1.0 µg pMD2.g packaging plasmids, and 10 µL TransIT-Lenti transfection reagent. On day 3, FBS was supplemented to 30%. On day 4, virus-containing media in each well was collected and replaced with DMEM supplemented to 30%

FBS. On day 5, virus-containing media was again collected and pooled with media collected the previous day. All supernatant was syringe filtered using 0.22 μm PES filters (Cytiva). The filtered virus was concentrated using the Lenti-X Concentrator (Takara Bio Inc.) and then stored in single use aliquots at -80°C.

**Lentiviral Overexpression and Localization in HeLa Cells**

To stably express viral constructs in HeLa cells, cells were seeded into a 10 cm plate in media supplemented with 8 μg/mL polybrene. Lentivirus was added to the media while cells remained in suspension and then plates were incubated overnight. At 24 h post-virus introduction, the media was replaced with normal culturing media. At 48 h post-virus introduction, cells were sorted for green fluorescence using a BD Melody fluorescence-activated cell sorter. Sorted cells stably expressing the viral constructs were seeded onto round glass coverslips in a 24-well plate format. The next day, cells were washed 2x with phosphate buffered saline (PBS) and fixed in a 4% final concentration of formaldehyde in PBS for 15 min.

*Cells expressing ectopic mCherry-tagged ETS constructs* (construct: ETS protein-mCherry-IRES-ZsGreen) were directly stained with DAPI (0.1 μg/mL) and washed 3x with PBS before mounting on glass microscope slides with ProLong Gold Antifade Mountant.

*Cells expressing ectopic GFP-tagged ETS constructs* (construct: GFP-β-Gal-fusions) or untagged ETV6 (construct: ETV6-IRES-ZsGreen) were washed 3x with PBS and permeabilized in 0.3% Triton-X 100 in PBS for 15 min. Permeabilized cells were washed 3x with PBS and then blocked at room temperature using 5% normal goat serum in PBS (blocking buffer) for 1 h. Cells were then washed 2x in a buffer containing 0.1% Tween-20, and 1% normal goat serum in PBS (PGST buffer). Respective primary antibodies were diluted in PGST to create the staining buffers. Cells expressing untagged ectopic ETV6 utilized an anti-ETV6 primary antibody (Sigma: HPA000264) diluted 1:250.

Cells expressing GFP-β-gal constructs required an anti-Vimentin primary antibody (diluted 1:100; Cell Signaling Technology: 5741S) as a cytoplasmic marker for immunofluorescence. Cells were incubated in staining buffer overnight at 4°C. The following day, cells were washed 3x with PGST and incubated with anti-rabbit IgG secondary antibody conjugated to Alexa Fluor 647 (Invitrogen: A21244) diluted 1:500 in PGST for 1 hr at 4°C. Cells were then stained with DAPI (0.1 µg/mL) and washed 3x with PBS before mounting on glass microscope slides with ProLong Gold Antifade Mountant.

**Confocal Imaging and Quantification of Immunofluorescence**

Cells mounted for localization were imaged with a Zeiss LSM 980 microscope. A 3 × 3 tiled image was taken using the Plan-Apochromat 40×/1.30 oil objective. Image processing was done with ImageJ and the Fiji image processing package. Using the Fiji image processing package, the DAPI channel was used to create nuclear regions of interest (ROIs) that were then used to measure the nuclear density of either the mCherry, GFP-β-gal, or Alexa Fluor 647 channel depending on the construct. The ZsGreen or Vimentin (Alexa Fluor 647) channels were then used to create cell ROIs. Pixels within the nuclear ROIs were subtracted from the cell ROIs to create cytoplasmic ROIs. These cytoplasmic ROIs were used to measure the cytoplasmic density of either the mCherry, GFP-β-Gal, or Alexa Fluor 647 channel, respectively. The densities in each nuclear ROI were averaged to obtain a nuclear density value and densities in each cytoplasmic ROI were averaged to give a cytoplasmic density value. The nuclear-to-cytoplasmic ratio was then calculated as the average nuclear density/average cytoplasmic density. At least 3 technical replicates were imaged for each construct and at least 50 cells were counted in each replicate as determined by the number of nuclear ROIs. All immunofluorescence images shown in this manuscript have been cropped and the brightness has been adjusted to allow for optimal visualization of protein localization.

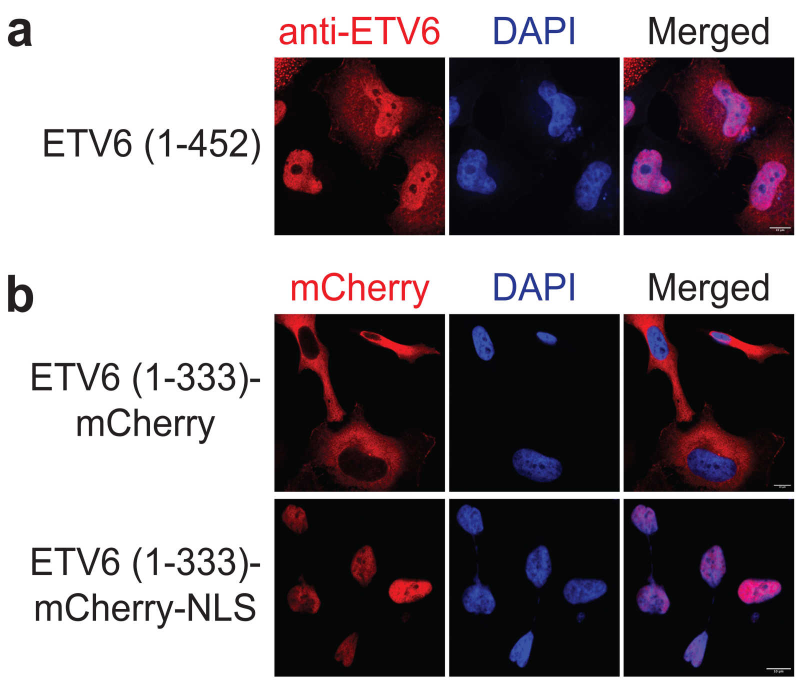

**Fig. S1. The ETV6 ETS domain is required for nuclear localization.**

a. ETV6 immunofluorescence microscopy z-stack images of HeLa cells bearing exogenous full-length, wild-type ETV6. Scale bar = 10 μm.

b. Fluorescence microscopy z-stack images of HeLa cells bearing the indicated exogenous ETV6 protein fragment. Scale bar = 10 μm.

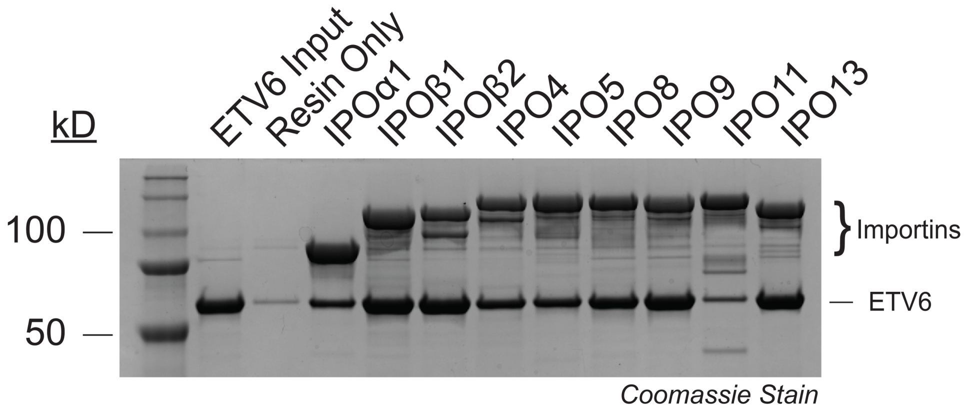

**Fig. S2. Full-length ETV6 exhibits an importin interaction specificity profile similar to that of the ETS domain.**

SDS-PAGE analysis of GST pull-down assays assessing the interaction between immobilized recombinant importin proteins (bait) and recombinant, full-length ETV6 (prey).

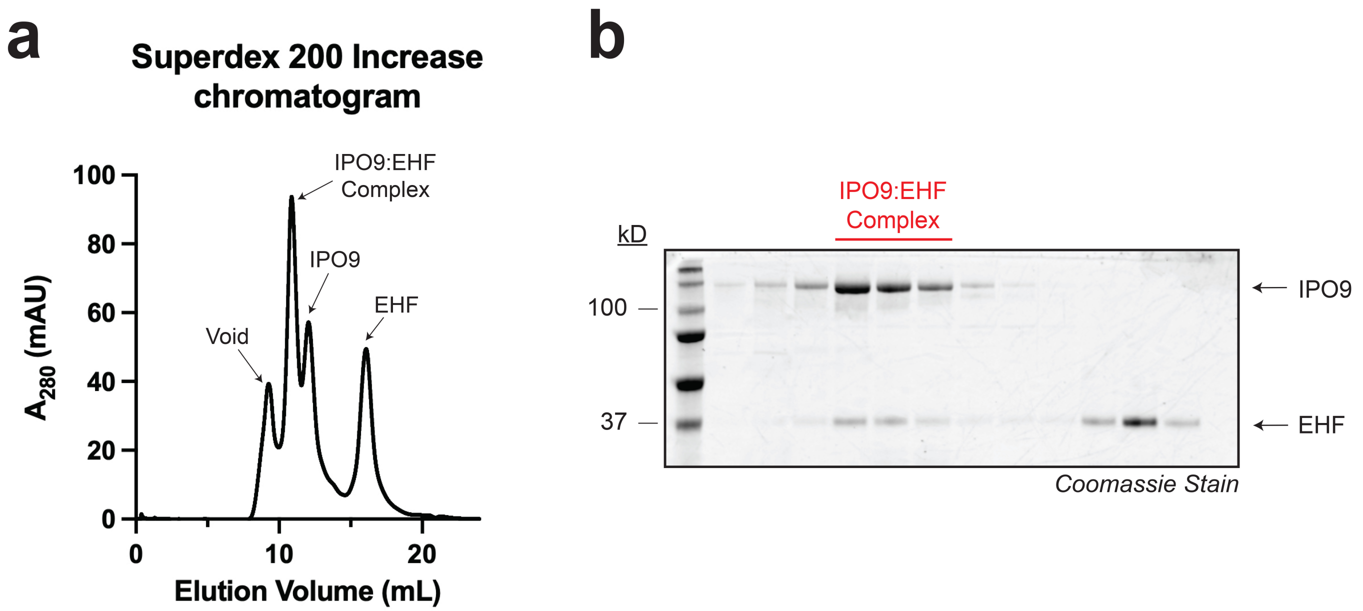

**Fig. S3. Purification of the IPO9:EHF complex via gel filtration chromatography.**

a. The gel filtration (Superdex 200 increase 10/300 GL) chromatogram obtained following injection of full-length, recombinant IPO9 and EHF mixed at a 1:5 molar ratio.

b. SDS-PAGE analysis of the gel filtration chromatography elution shown in panel (a). Lanes correspond to fractions ranging from elution volume 8-18 mL.

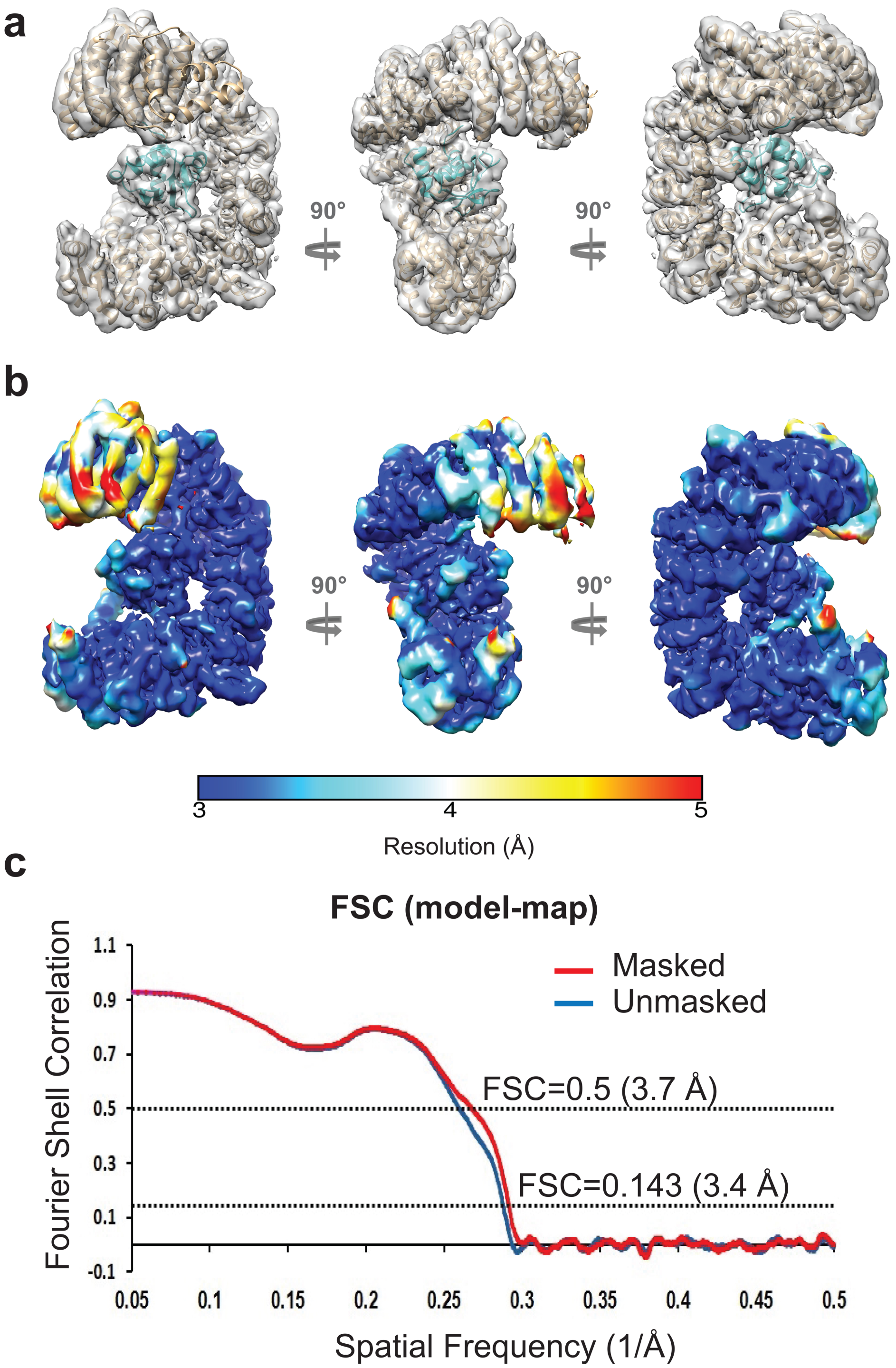

**Fig. S4. Cryo-EM electron density map quality and model validation for the IPO9:EHF complex.**

a. The cryo-EM density map with the IPO9:EHF model superimposed.

b. Cryo-EM density map colored by local resolution.

c. Fourier shell correlation (FSC) analysis of the IPO9:EHF cryo-EM map and atomic model.

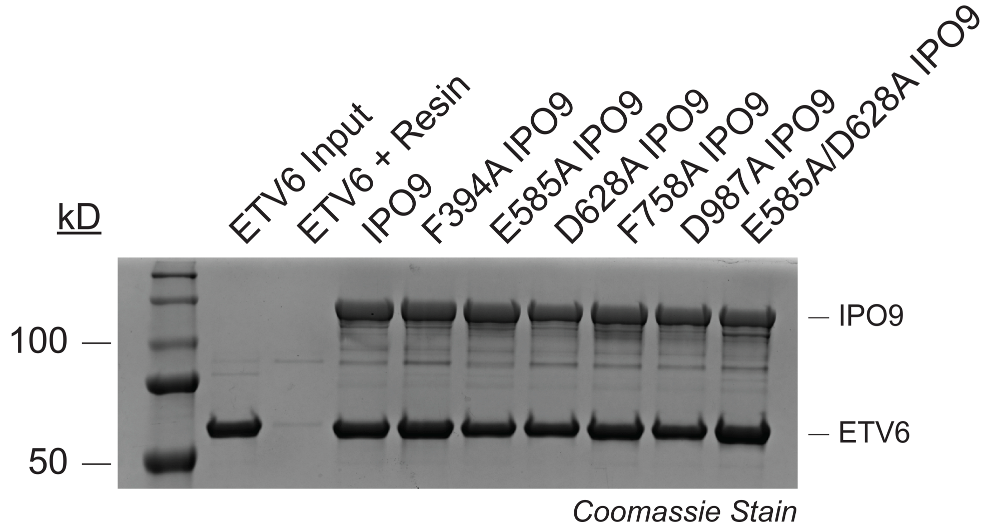

**Fig. S5. Single and double IPO9 point mutations have no detectable effect on IPO9:ETV6 complex formation.**

SDS-PAGE analysis of GST pull-down assays assessing the interaction between immobilized recombinant IPO9 protein variants (bait) and full-length, recombinant ETV6 (prey).

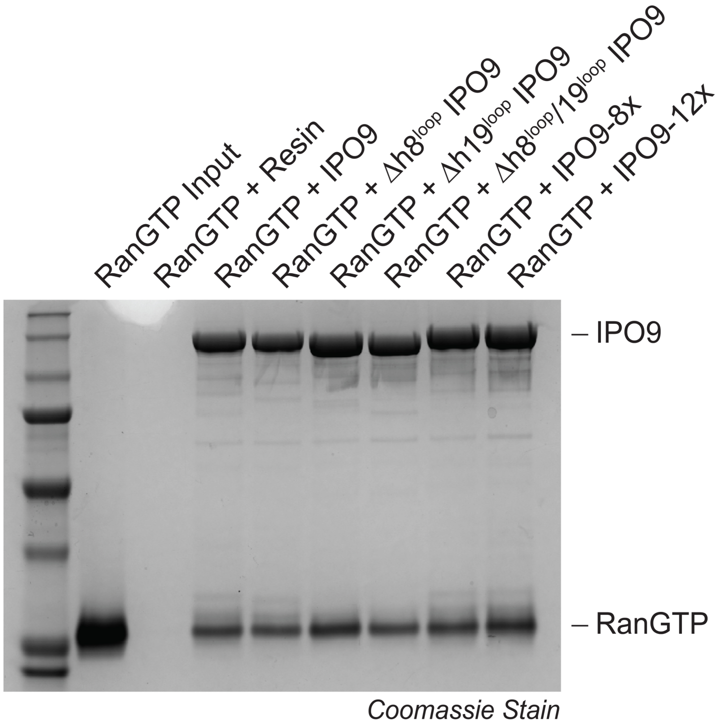

**Fig. S6. IPO9 batch mutations and loop substitution variants differentially affect EHF and RanGTP binding.**

SDS-PAGE analysis of GST pull-down assays assessing the interaction between immobilized recombinant IPO9 protein variants (bait) and recombinant RanGTP (prey).

**a**

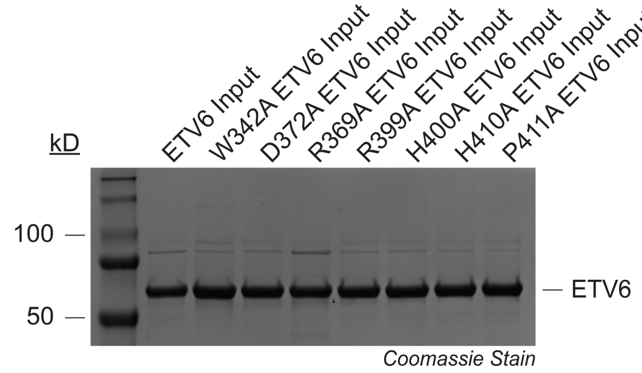

**b**

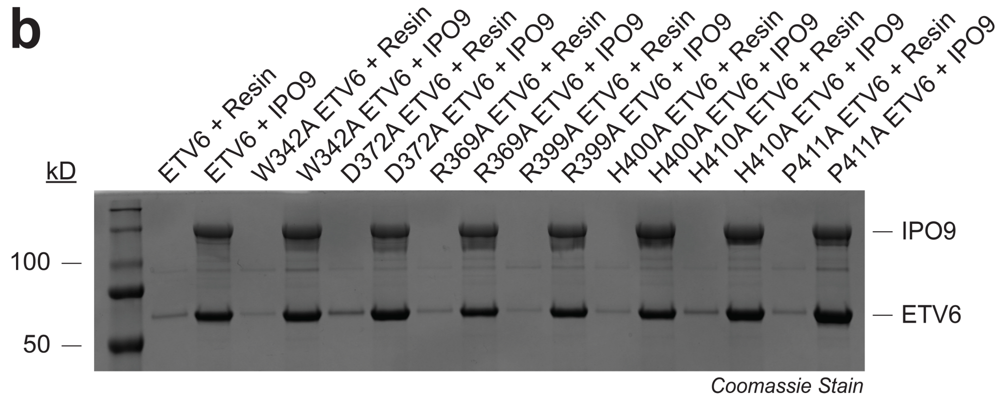

**Fig. S7. Single ETV6 point mutations have no detectable effect on IPO9:ETV6 complex formation.**

a. SDS-PAGE characterization of ETV6 protein variants to confirm purity and concentration.
b. SDS-PAGE analysis of GST pull-down assays assessing the interaction between immobilized recombinant GST-IPO9 (bait) and full-length, recombinant ETV6 protein variants (prey).

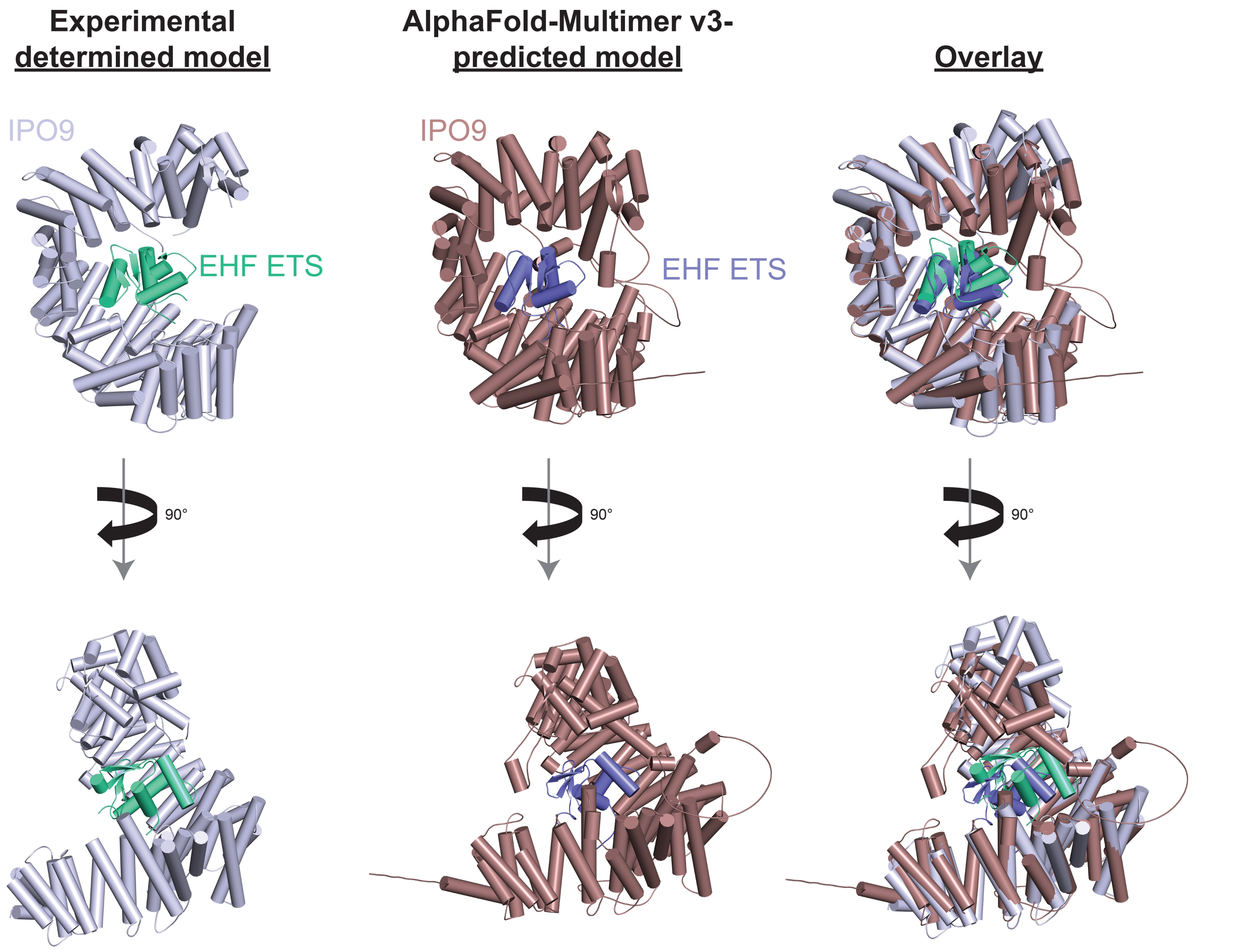

**Fig. S8. A comparison of the experimentally determined IPO9:EHF complex structure and AlphaFold-Multimer v3-predicted model.**

Model comparisons reveal key differences in IPO9 conformation and EHF orientation relative to the concave IPO9 surface. Consequently, the AlphaFold-Multimer v3-predicted model fails to identify several critical interface features, including interaction hotspots on the IPO9 concave surface and the h19$^{loop}$.

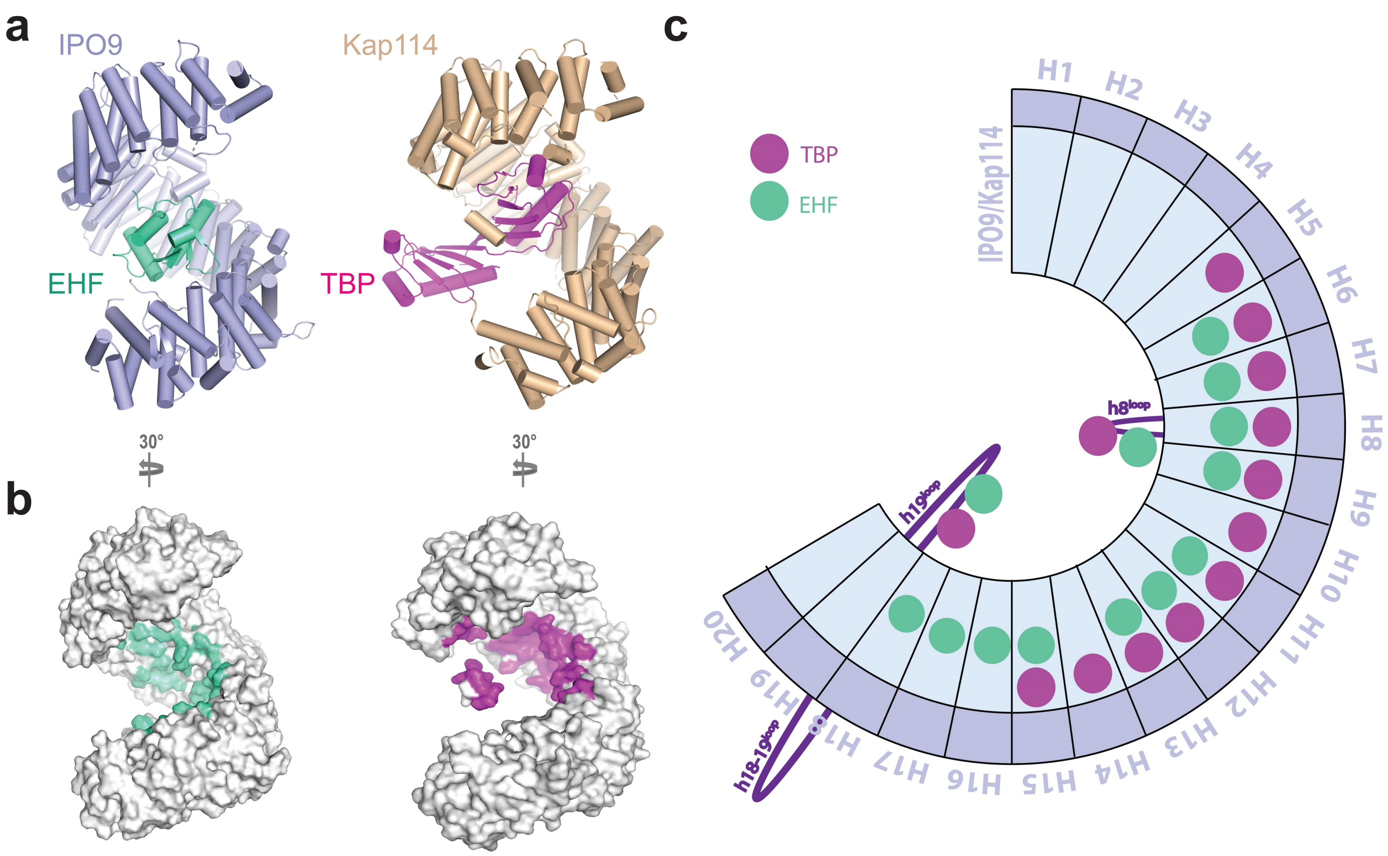

**Fig. S9. Structural comparisons of IPO9:EHF and Kap114p:yTBP complexes reveal distinct cargo recognition mechanisms.**

a. Side-by-side structural comparison of the IPO9:EHF and Kap114p:yTBP complexes.
b. Surface representations of IPO9 and Kap114p with their cargo contact interfaces highlighted (EHF in green and yTBP in magenta, respectively).
c. Schematic alignment of HEAT repeats in IPO9 and Kap114p, with HEAT repeat that engage EHF or yTBP indicated (colored circles).

**Table S1. Statistics of Cryo-EM data collection, processing, and refinement for IPO9:EHF.**

| | IMP9-EHF |
|---|---|
| **Data collection and processing** | |
| Magnification | 105,000 |
| Voltage (kV) | 300 |
| Electron exposure ($e^{–}/Å^2$) | 60 |
| Defocus range (μm) | -0.9 to -2.2 |
| Pixel size (Å) | 0.41 |
| Symmetry imposed | C1 |
| Initial particle images (no.) | 3,296,171 |
| Final particle images (no.) | 136,260 |
| Map resolution (Å) | 3.48 |
| FSC threshold | 0.143 |
| **Refinement** | |
| Initial model used (PDBID) | AF-Q9NZC4-F1-v6; 6N1Z |
| Model resolution (Å) | 3.2/3.4/3.7 |
| FSC threshold | 0/0.143/0.5 |
| Map sharpening B factor ($Å^2$) | -148 |
| Model composition | |
| Non-hydrogen atoms | 16241 |
| Protein residues | 1022 |
| R.m.s. deviations | |
| Bond lengths (Å) | 0.004 (1) |
| Bond angles (°) | 0.908 (0) |
| **Validation** | |
| MolProbity score | 1.28 |
| Clashscore | 5.18 |
| Poor rotamers (%) | 0.00 |
| Ramachandran plot | |
| Favored (%) | 98.21 |
| Allowed (%) | 1.79 |
| Disallowed (%) | 0.00 |

**Supporting Information References**

17. Anonymous, Schrödinger, LLC. The PyMOL Molecular Graphics System. *Version 2.5.5. Schrödinger, LLC*.
